# Attentional selection is a neuroeconomic decision

**DOI:** 10.64898/2026.05.22.727072

**Authors:** Christoph Strauch, Stefan Van der Stigchel, Sanyukta Nair, Damian Koevoet

## Abstract

Sensorimotor decisions require balancing extrinsic rewards against intrinsic action costs. How these signals are integrated during rapid eye-movement choices remains unclear. We show that human saccade selection follows cost-benefit computations consistent with neuroeconomic decision-making. Across two experiments, participants maximized rewards and minimized effort-costs. Reward titration revealed that more effortful saccades required higher monetary rewards to be chosen equally often. While effort-costs affected choices linearly, small reward differences shifted choices more than equally-sized larger ones. Furthermore, effort-cost (but not reward) differences regulated decision engagement: Deliberation was extended only when cost differences were large. Together, costs and rewards play dissociable roles in saccade selection, positioning eye movements as a tractable model system for economic decision-making.

## Main

Humans must select what to attend and what to ignore multiple times per second through rapid eye movements (’saccades’) and covert shifts of attention. Classical accounts explain visual selection in terms of priority signals arising from bottom-up salience, top-down goals [1], and more recently, selection history [2, 3]. These frameworks specify which stimuli are relevant, but ignore whether intrinsic costs of shifting attention constrain the resulting choice. In contrast, research in motor control, decision science, and economics that have long emphasized that action selection reflects trade-offs between expected rewards and intrinsic costs [4–8].

It is well established that expected reward is encoded in saccadic choices and that saccades are made proportionally to relative reward rates [9, 10]. Work in motor neuroscience has further demonstrated that saccade movement trajectories encode the value of selected options [5, 11, 12]. At the more upstream stage of selection itself, utility maximization, rather than mere reward maximization, has been argued to be central [13, 14]. Full economic arbitration at the point of selection, however, would necessitate that costs and rewards jointly shape which target is committed to, not only the vigor with which it is pursued. Substantiating an economic account of attentional selection further requires that both rewards and costs can be measured independently.

Indeed, recent work suggests the minimization of intrinsic effort-costs of attentional shifts to be a fundamental driver of attentional selection [15–19] (see [20] for a review and other types of saccade costs), with extra invested effort changing kinematics [21]. However, such costs are typically assumed and invoked [13, 14] or inferred from behavioral residuals [e.g. 16] rather than measured independently from the selection behavior they seek to predict. We recently used pupil dilation - a well-established marker of cognitive and motor effort [22–26] - to quantify the effort-costs associated with saccade planning [17, 18]. These physiologically-measured costs accurately predicted saccade choices (affordable over costly ones) [18], even when bottom-up salience differed between targets [19].

Saccade selection is thus driven by reward maximization and effort minimization when studied in isolation. However, it is unknown how rewards and effort-costs are integrated during saccade selection. We here hypothesized that the oculomotor system integrates rewards and effort-costs following principles of economic decision-making. Put simply, the utility of attending to a target can be thought of as reward minus cost. However, economic decision-making can take different forms and levels of sophistication. We thus formulated three increasingly stringent criteria of economic decision-making to test whether saccade selection follows neuroeconomic principles:

1. Costs and rewards are integrated on a common utility scale (rewards - costs), as formalized in expected utility theory [6] and widely applied to effort-based decisions [5, 27, 28]. Under this view, costs and rewards should trade off, such that more costly options require higher rewards to be chosen equally often.
2. More structured economic integration implies that costs and rewards act as distinct signals rather than one undifferentiated signal [4].
3. Finally, if choices reflect utility comparisons, decision time should scale with utility differences as proposed by sequential sampling models [29, 30] and neural evidence [31].

Our results show that saccade selection fulfills all three criteria, indicating elaborate economic decision-making at the level of saccades: higher-cost saccades were chosen when offset by greater rewards, costs and rewards influenced choices differently, and decision engagement only scaled with cost but not reward differences. These data demonstrate that neuroeconomic principles operate at the sensorimotor level of eye-movement selection.

## Results

### Saccade preferences differ across directions

In Experiment 1, twenty human participants freely selected one of two targets in a saccade choice task. Saccade targets differed in shape which signaled its associated reward (0, 1 or 50 points) that would later translate into monetary rewards. Shapes and their positions were fully orthogonal (see Figure 1a). In Experiment 2, thirty-two human participants performed a similar baseline block without reward differences. In a second block, rewards were dynamically updated depending on direction. The amount of points associated with a specific shape was presented at the start of each trial (see Figure 2a). Initially, all directions had a reward value of 100. Rewards were then updated based on previous choices using a dynamic ranking algorithm inspired by chess ratings [ELO, see 32]. Choices that matched expected outcomes led to smaller reward updates (see Figure 2b).

**Figure 1:**
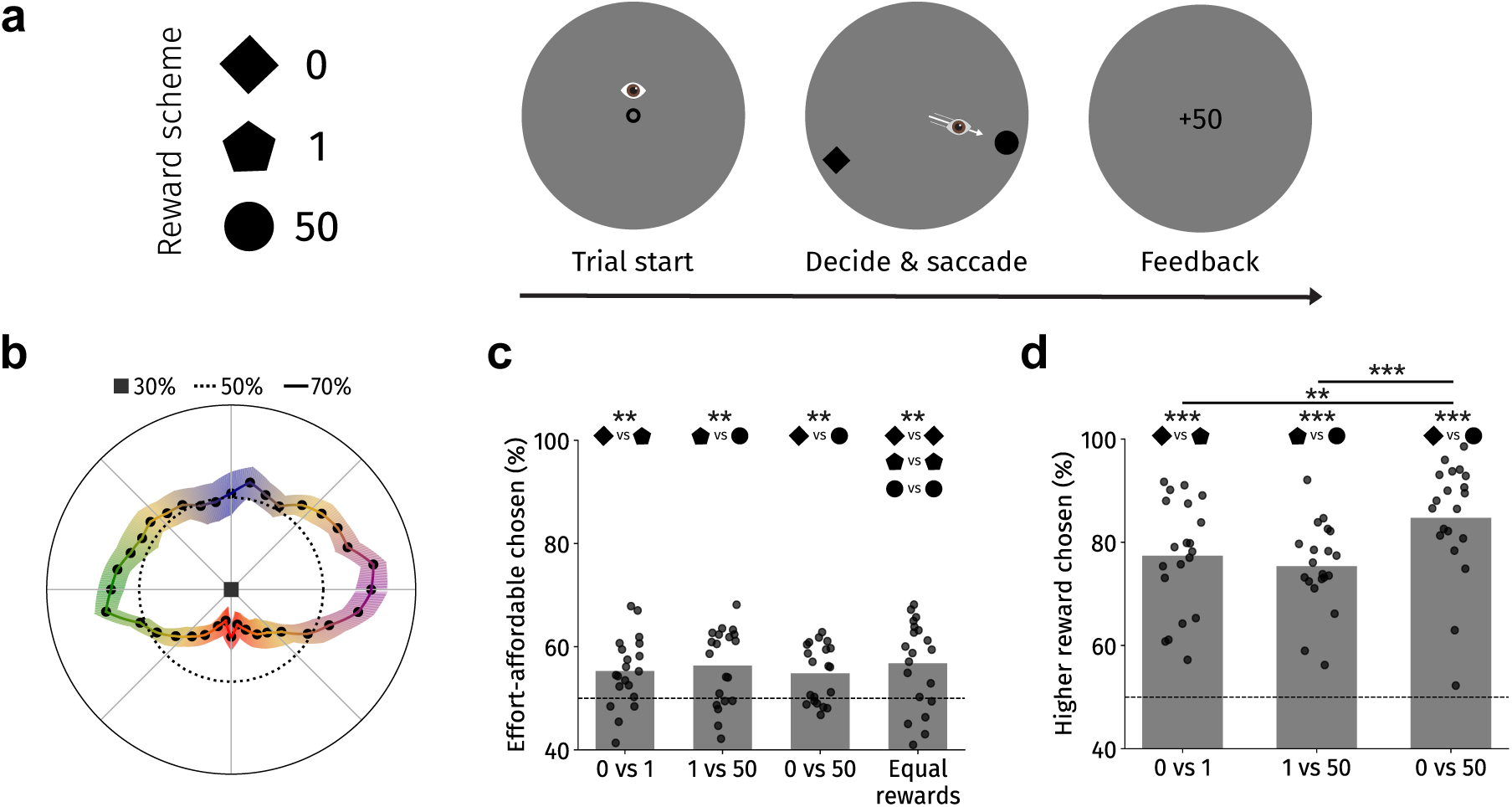
Saccade choices are determined by effort-costs and rewards (Experiment 1). **a** Participants choose between two targets by saccading; reward associated with each shape is instructed and feedback is provided after each trial. **b** Saccade preferences varied across directions. **c** Across reward combinations, lower effort-cost (affordable) targets were preferred over costlier targets. **d** Preferences for the higher-reward target did not differ between 0 vs 1 and 1 vs 50 point differences, and were strongest for 0 vs 50 points. **: *p* < 0.01, ***: *p* < 0.001.

**Figure 2:**
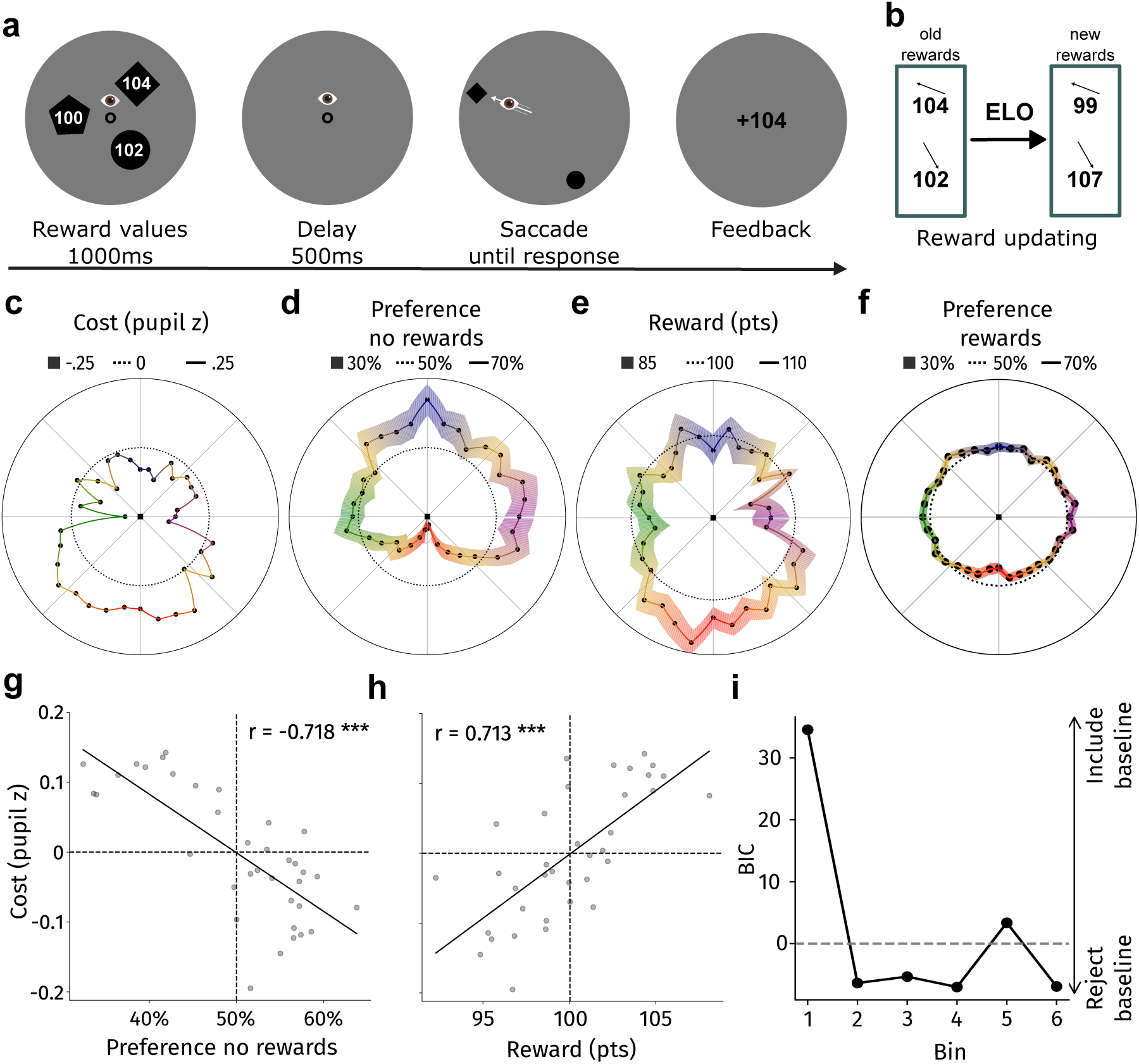
Effort-costs predict preferences and are offset by titrated rewards (Experiment 2). **a** Dynamic reward block: Participants first see three shapes containing reward information. Participants then select either of two presented shapes. Feedback signals the obtained reward. **b** Rewards are updated online. **c** Out-of-sample pupil-inferred saccade costs [18]. **d** Baseline preferences without rewards varied across directions. **e** Final rewards were higher for less preferred baseline directions. **f** Affected by both costs and rewards, preferences vanished almost completely. **g** Cost minimization predicts preferences without rewards. **h** Costs were linked to final dynamically updated rewards, effectively translating physiological into economic units. **i** BIC of a LME predicting preferences from no reward (baseline) preferences and rewards: baseline preferences became increasingly less predictive as rewards were dynamically updated. *** *p* < 0.001

We first examined participants’ saccade preferences across directions using trials from Experiment 1 (Figure 1b) and baseline trials (no reward difference) from Experiment 2 (Figure 2d; total *N* = 52). In line with previous work [18, 33–36], we found saccade preference asymmetries across directions. Specifically, we observed a preference for cardinal over oblique targets and for upward over downward targets (*β*_obliqueness_ = 0.035, *z* = 3.144, *p* = 0.002; *β*_verticalness_ = 0.041, *z* = 4.546, *p* < 0.001; no difference between left and right sides of the screen, *β*_horizontalness_ = 0.009, *z* = 1.089, *p* = 0.276).

### Saccade costs and rewards predict saccade preferences

Saccade costs were independently established by measuring pupil size, an established marker of effort [22–25], during saccade planning in a different group of participants [18]. As saccade costs differ across directions [16–18], this allowed us to test whether saccade cost minimization predicts saccade selection. For each trial, we predicted participants to choose the less effortful saccade target, and found that saccade costs were minimized in all conditions in Experiment 1 (Figure 1c; all *p*s < 0.01).

Turning to reward maximization, we found that participants preferred saccading to targets associated with higher rewards in all conditions in Experiment 1 (Figure 1d; *p*s < 0.001). The preference for the more rewarding saccade target was more pronounced in the 0 vs 50 points condition that featured the biggest reward difference, compared with the 1 vs 50 points condition (*t*(19) = −5.702, *p* < 0.001, *d* = −1.275) and the 0 vs 1 point condition (*t*(19) = −3.852, *p* < 0.001, *d* = −0.861; with no further differences between conditions).

We again observed cost minimization and reward maximization in Experiment 2 when reward differences were more continuous. Participants preferred more affordable targets over costlier ones (Δcost: *β* = 1.206, *z* = 5.192, *p* < .001) and more rewarding targets over less rewarding ones (Δreward: *β* = 0.053, *z* = 9.026, *p* < .001). There was no interaction between cost and reward differences (Δcost : Δreward: *β* = 0.003, *z* = 0.520, *p* = .603).

### A cost-reward tradeoff predicts saccade selection

Having established saccade cost minimization and reward maximization in both experiments, we tested whether these factors are dynamically traded-off following neuroeconomic principles (i.e., a common utility scale, first principle formulated above). For example, higher cost targets may be selected only when the associated reward is large and vice versa.

To provide first insight into this question, we split trials based on whether participants selected the more affordable or the costlier target. We reasoned that participants should obtain higher reward on average when picking the costlier target to make their effort-cost investment worthwhile (see Figure 3a for Experiment 1 and Figure 3b for Experiment 2). Indeed, we found that participants on average required higher reward to choose more costly targets (Experiment 1: *t*(19) = 4.184, *p* < 0.001, *d* = 0.936; Experiment 2: *t*(31) = 4.851, *p* < 0.001, *d* = 0.858). These data indicate an economic trade-off between (physiologically-inferred) costs and (monetary) rewards during saccade selection. We leveraged the more continuous differences in reward during the main block of Experiment 2 to examine this trade-off in more detail. When selected targets were more affordable, they were associated with lower rewards (Figure 3c, *r* = −0.288, *p* < 0.001); costlier selected targets were associated with higher rewards (Figure 3d, *r* = 0.312, *p* < 0.001). These links scaled monotonically with cost differences.

**Figure 3:**
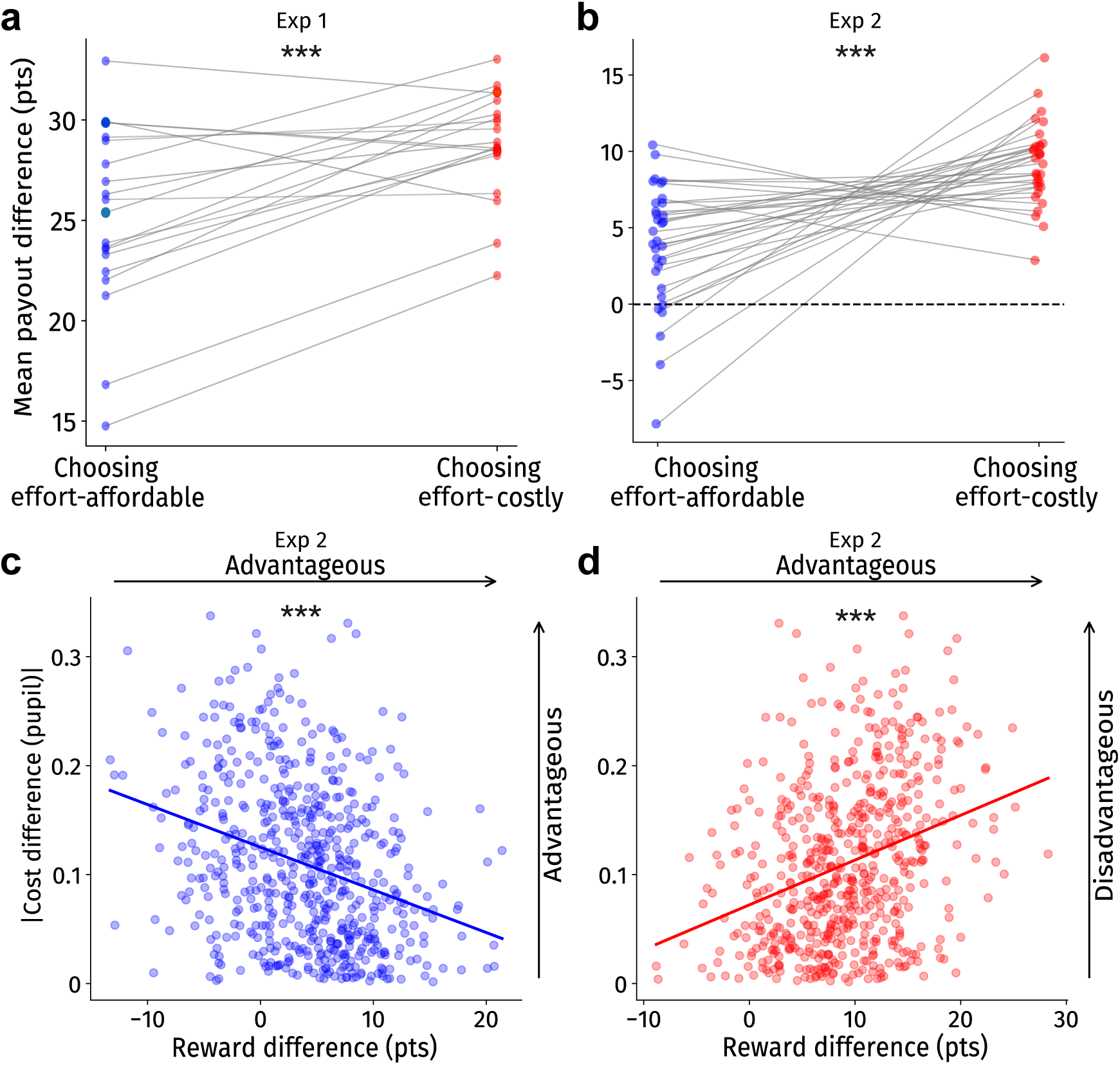
Costlier saccades require greater reward. **a,b** Participants accept lower rewards for the lower-cost target (blue) and require higher rewards for the costlier target (red). **a** for Experiment 1, **b** for Experiment 2. Dots denote participant means. **c** For selections of the lower-cost target (blue), the minimum reward participants accepted decreased monotonically as the cost advantage increased, reflecting greater willingness to choose less rewarding but cheaper options (Experiment 2). **d** For selections of the higher-cost target (red), the minimum reward required increased with cost, reflecting compensation for choosing the more effortful target. c,d: Dots denote pooled means for unique target combinations (Experiment 2). *** *p* < 0.001

### Extrinsic rewards compensate for intrinsic costs

Extending this trade-off further, we next asked whether intrinsic costs can be offset by extrinsic rewards (Experiment 2, main block). To this end, we titrated rewards associated with specific directions with the aim of allowing fine-grained compensation of costs through rewards that should ultimately result in a uniform saccade preference map. This would indicate that extrinsic rewards could compensate for intrinsic cost differences across directions.

Following reward updates, the baseline direction preferences (Figure 2d) largely disappeared (Figure 2f), producing a near-uniform saccade preferences map across directions. If truly trading off intrinsic costs against extrinsic (monetary) reward, then the payoffs across directions should directly mirror the intrinsic saccade costs map established using pupil size. Indeed, the reward titration map paralleled the cost map (*r*_(rewards, costs)_(34) = 0.713, *p* < 0.001, Figure 2h), establishing a cross-unit mapping between intrinsic costs and extrinsic rewards. This demonstrates that fine-grained intrinsic cost differences are (implicitly) accessible during saccade selection, and costs can be compensated through economic incentives. These data provide evidence that intrinsic costs and rewards are directly traded off against each other, satisfying our first criterion of neuroeconomic decision-making.

### Costs and rewards map differently onto choices

Costs and rewards are thought to reflect distinct signals that differ in terms of computation and neural underpinnings in substantially more explicit tasks [4, 37]. Having established that saccade selection acts as a neuroeconomic process balancing costs and rewards, we therefore asked *how* costs and rewards map onto choices.

To reveal the mapping of costs onto choices, we compared cost differences per target combination with the probability of choosing either target. We observed a linear mapping of costs onto choice: higher costs resulted in reduced preference (Experiment 1 Figure 4a; Experiment 2 Figure 4b). This relationship vanished once costs were compensated by rewards and thus absorbed into them via adaptive reward titration (Figure 4c). In contrast, reward differences mapped nonlinearly onto choice probability: larger reward differences exerted a diminishing marginal influence on choice. This mapping was well-described by a psychometric function with a symmetric lapse parameter, as commonly observed in perceptual decision-making (*R*^2^ = 0.993 across 15 bins, [consistent with 38]). This sigmoidal mapping is consistent with classical accounts of diminishing returns [39]. Thus, while saccade costs linearly mapped onto saccade choices, rewards affected choices through a sigmoidal mapping. This strongly indicates that saccade costs and rewards drive saccade selection in distinct ways, fulfilling our second criterion of neuroeconomic decision-making.

**Figure 4:**
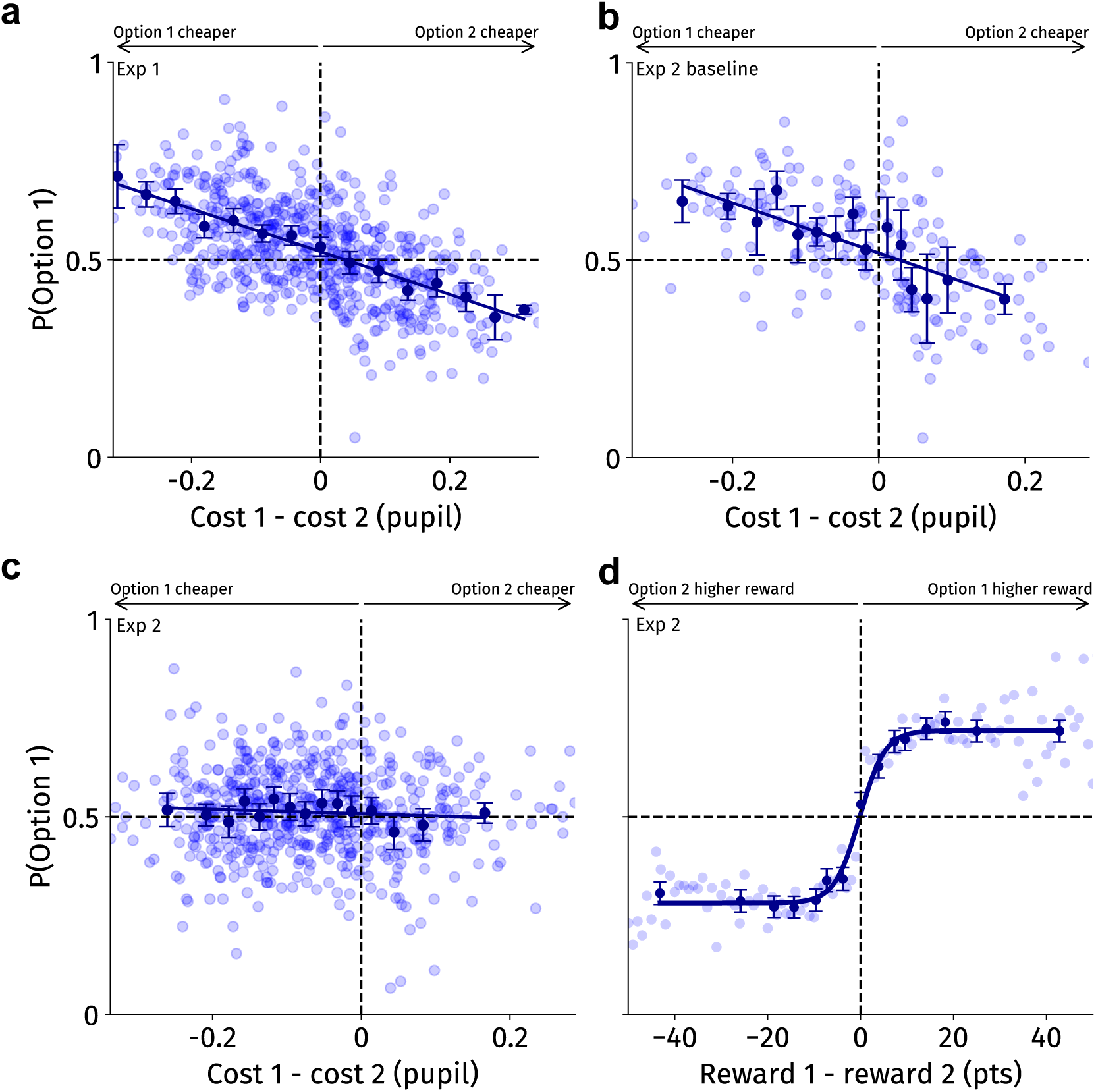
Costs and rewards map differently onto choice (Experiments 1 and 2). **a** (Experiment 1) and **b** (Experiment 2 baseline) choices followed linear cost minimization. **c** Reward titration absorbed effort-costs in Experiment 2 (after bin 1). **d** Choice probability varied sigmoidally with reward differences: small reward differences influenced choices disproportionally strong, whereas larger differences yielded diminishing marginal effects. **a-c** Dots reflect cost differences between target combinations (x-axis) with the respective proportion of choices for the lower effort-cost option (y-axis) across all trials and participants. **d** Dots reflect unique reward differences (x-axis) with the respective proportion of choices for the more rewarding option (y-axis). Bold dots reflect bin means of equal trial number in all panels alongside 95% confidence intervals.

### Larger utility differences speed up decisions

If saccade choices reflect an even more stringent form of economic decision-making, trials with larger utility differences should be associated with faster response times. That is, more complex and elaborate decision-making should take longer [4, 29, 40]. We indeed found shorter saccade latencies when rewards differed in Experiment 1 (compared with no reward difference), likely reflecting a larger utility difference between the two offered targets, given how predictive reward was of choices (*t*(19) = 4.308, *p* < 0.001, *d* = 0.963; Figure 5a). In contrast, saccade latencies were longer under the ELO regime relative to the no-reward baseline in Experiment 2 (*t*(31) = 3.605, *p* = 0.001, *d* = 0.637; Figure 5b). This paradox resolves once effective utility (rewards - costs), not nominal reward, is treated as the driver of response times. Under the reward titration regime, the utility of all potential saccade targets is effectively equated as evidenced by the near-uniform saccade preference map (Figure 2f). Therefore, utility differences were larger in the baseline block compared with the later titration blocks. We thus found a pattern that is consistent with decision-making models [4, 29, 40]: Saccade choices with larger utility differences require more elaborate, or slower, decision-making. This also challenges the common assumption that rewards speed up saccades [41]. When rewards are titrated to equalize utility across options, they can reduce rather than increase utility differences, making choices harder and slowing responses instead.

**Figure 5:**
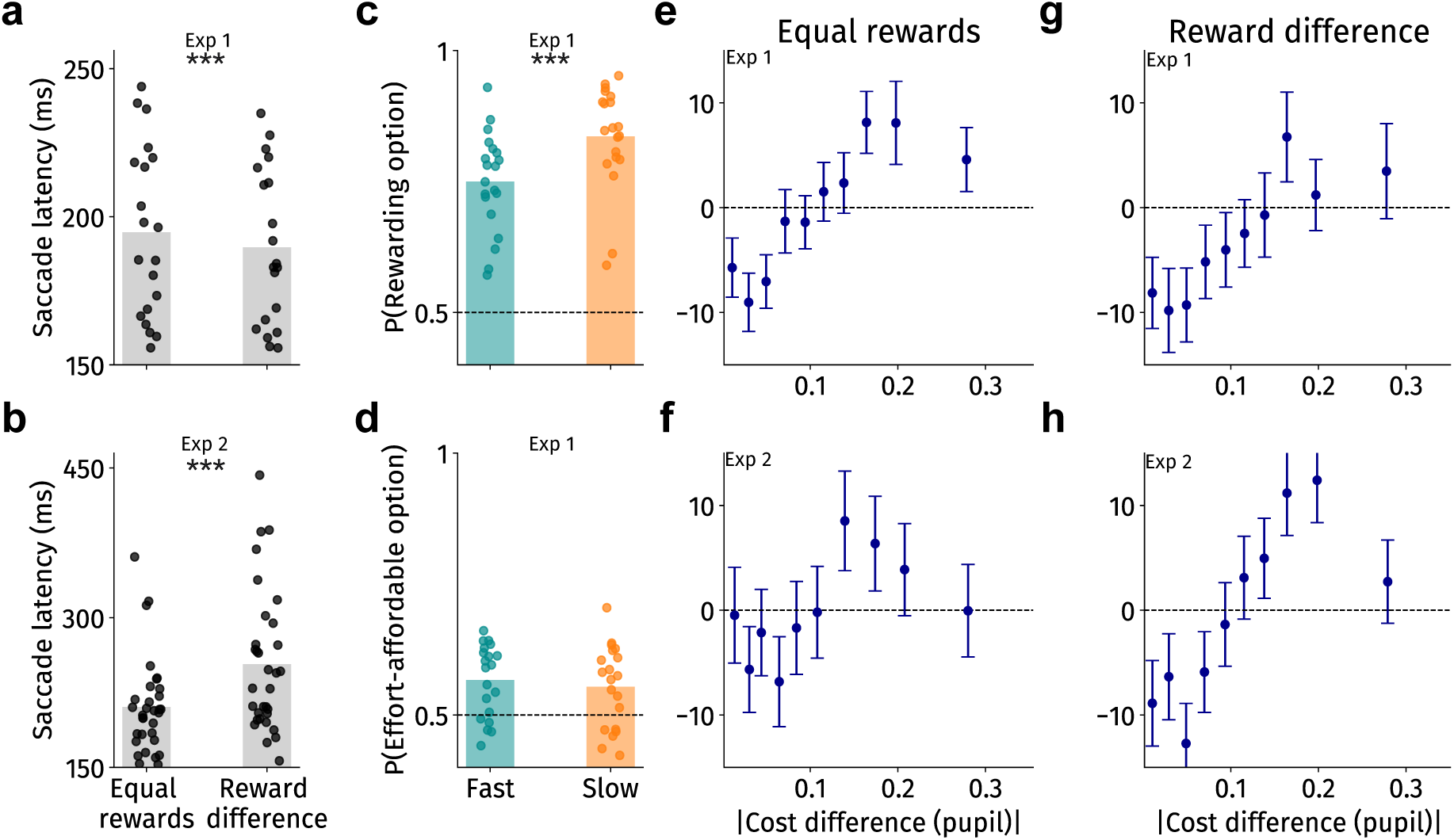
Costs regulate decision depth, rewards gain influence with increasing decision time (Experiments 1 and 2). **a** Average saccade latencies in Experiment 1 accelerated through reward differences (bigger utility gap). **b** Average saccade latencies in Experiment 2 slowed when reward differences were present (smaller utility gap). **c,d** Longer decisions in Experiment 1 increased selection of higher-reward, but not lower-cost, targets (median split); dots denote participant means. With the minimized utility differences in Experiment 2, neither costs nor rewards predicted saccade selection better with increased decision times. **e-h** Demeaned saccade latencies (per participant and location, for visualization only) are shortest for choices involving small cost differences and increase nonlinearly as cost differences grow before decreasing in turn. Errorbars denote 95% confidence intervals.

### Slow decisions allow more reward-optimal eye movements

Longer and thus plausibly more elaborate decisions should therefore be associated with higher choice quality, i.e., higher utility yield than faster ones. We thus median-split trials into slow and fast decisions for Experiment 1. We found choices to be more reward-driven when taking more time to decide (*t*(19) = 8.155, *d* = 1.823, *p* < 0.001, see Figure 5c). In contrast, costs were not differently predictive for slower compared with faster decisions (*t*(19) = −1.150, *d* = −0.257, *p* = 0.264, see Figure 5d). This may reflect costs biasing choices inherently whereas rewards had to be inferred from perception and memory, hence benefiting from additional time. These data indicate that saccade costs and reward operate on different timescales: While cost differences affect selection regardless of decision time, explicit reward differences require time to guide selection.

In Experiment 2, utility differences between targets were minimized by design, providing a situation where deliberation cannot or only minimally improve choice quality. Therefore, we expected that the modulatory role of decision speed on reward-driven selection should be reduced or absent when titrating rewards. Consistent with this, we observed no reliable interactions between the reward difference of the two options and saccade latency (Δreward(z) ∗ latency: *β* = 0.039, *z* = 1.852, *p* = 0.064), nor between cost differences and latency (Δcost(z) ∗ latency *β* = −0.006, *z* = −0.343, *p* = 0.732). Together, longer decisions allowed more utility-optimal choices only when utility differences between options were large enough (i.e., in Experiment 1). In these cases, increased optimality reflected a shift toward reward-driven choice rather than increased cost minimization.

### Cost differences regulate decision engagement

Until now, we have considered saccade selection as the integrative outcome of cost-reward trade-offs. However, the decision to integrate these determinants of saccade selection itself must also be associated with effort. Therefore, a truly cost-optimal decision makes an intriguing meta-economic prediction that reaches beyond the initially defined three criteria: if utility (i.e., costs and reward) differences between decision options are very small, it may be more optimal to choose randomly rather than forming an elaborate decision. Specifically, the potential gains of a specific choice may not always exceed the effort of forming an elaborate decision. If saccade selection is resource-optimal, engagement should be regulated such that a more elaborate decision-making process is recruited only when the expected value of deciding exceeds the expected value of random choice [4, 14, 42].

To infer decision time, we modeled saccade latency as a function of cost difference (linear and quadratic terms) while taking differences in latency across direction and participant into account. Costs consistently showed a curvilinear relationship with decision times: Larger, and especially intermediate, cost differences were associated with longer saccade latencies in conditions without reward differences (Experiment 1 Δcost: *β* = 200.8, *z* = 7.61, *p* < 0.001; Δcost^2^: *β* = −457.9, *z* = −4.82, *p* < 0.001; Figure 5e; baseline Experiment 2 Δcost: *β* = 170.5, *z* = 4.55, *p* < 0.001; Δcost^2^: *β* = −415.1, *z* = −3.13, *p* = 0.002; Figure 5f). The same pattern held when rewards differed (Experiment 1 Δcost: *β* = 128.9, *z* = 5.43, *p* < 0.001; Δcost^2^: *β* = −232.8, *z* = −2.78, *p* = 0.005; Figure 5g; Experiment 2 Δcost: *β* = 244.0, *z* = 8.17, *p* < 0.001; Δcost^2^: *β* = −524.2, *z* = −4.85, *p* < 0.001; Figure 5h), whereas reward differences showed weak and inconsistent effects (Experiment 1: *β* = −0.045, *z* = −2.01, *p* = 0.045; Experiment 2: *β* = 0.018, *z* = 0.35, *p* = 0.727). Reward differences assessed using categorical contrasts did not consistently predict decision times (Experiment 1: both contrasts *z* < −1.20, both *p* > 0.24, see Supplementary Figure 1a; Experiment 2: Δreward: *z* = −0.22, *p* = 0.824; Δreward^2^: *z* = 0.25, *p* = 0.807, see Supplementary Figure 1b).

The link between large effort-cost differences and increasing latencies is in line with decisions becoming easier as differences between options grow and uncertainty decreases [e.g. 29, 40]. Together, this implies a regulating role of costs in saccade selection, such that cost differences determine the degree of elaboration between targets: smallest cost differences lead to decision disengagement, intermediate ones require more elaborate and difficult decision-making, and largest differences produce easy choices.

## Discussion

We demonstrated that attentional selection operates as a neuroeconomic decision process that incorporates distinct measurable cost and reward signals. Both independently and physiologically-indexed saccade costs [18] and monetary rewards predicted saccade decisions. Participants precisely traded off intrinsic saccade costs with monetary rewards. We further found linear cost minimization and reward maximization that was characterized by diminishing returns (i.e., small reward differences being almost as important as very large reward differences). Saccade latencies provided further evidence for advanced economic arbitration in saccade selection, as decisions with large utility differences were reached faster than decisions with small utility differences. Further, saccade latencies revealed a meta-economic effect: decisions were more elaborate when intrinsic effort-cost differences were larger, likely offsetting the cost of deeper deliberation. More elaborate decisions, in turn, were associated with more reward-optimal behavior. Together, saccade costs and rewards are distinct drivers of saccade selection that differently map onto decisions, and act on different temporal scales. The integration of costs and rewards determines the outcome of saccadic decisions (see Figure 6 for a testable economic decision architecture based on the present results). This puts forth saccade selection as low-level human economic decisions, providing a tractable model for economic decision-making more generally.

**Figure 6:**
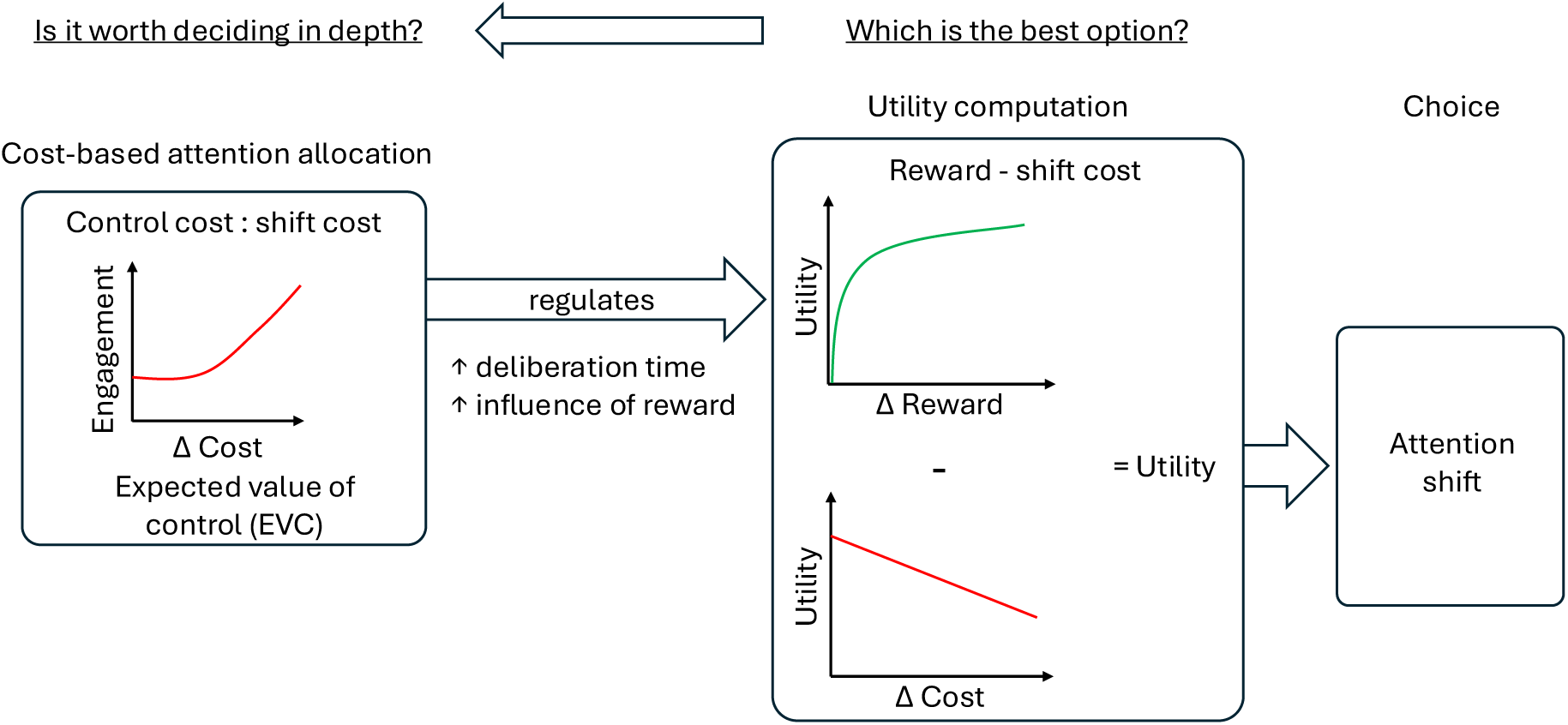
Attentional selection as an economic decision that seeks to select optimally. This entails a utility computation from rewards and costs that exert their influence differently (i.e., sigmoidally vs linearly). Net utility determines the attentional choice (such as a saccade). If cost differences exceed control costs, decisions are made more elaborately, as demonstrated by longer decision times and an increased influence of rewards on choices. Large utility differences go in hand with faster choices.

We replicated a saccade preference toward targets with lower effort-costs [18, 19] in trials without reward difference in both experiments. Extending this, we found that saccade cost remained predictive of saccade selection when options differed in reward. Saccade costs and monetary rewards were dynamically traded off: participants chose less rewarding targets if they were able to pick a low-cost target and vice versa. We detailed this trade-off in Experiment 2 by titrating the reward associated with each direction through an ELO procedure. We found that (implicit) cost information is accessed with high fidelity and traded off against explicit monetary reward to a degree that saccade preferences across directions vanished almost completely. The resulting map of rewards per direction should thus reflect latent saccade costs, which was validated by demonstrating a correlation with the previously established pupil-size indexed cost map [17, 18]. With access to this reward-inferred costs map, one can translate physiologically-inferred costs into monetary rewards and vice versa. Besides externally validating the claim of effort-linked pupil dilation as a useful metric of saccade cost [17, 18, 20], this lays the groundwork for computational models on a common scale. Such modeling in turn allows for quantitative predictions about decision thresholds or engagement, for instance.

Our work is not necessarily bound to monetary rewards. Perceptual information could itself behave like a reward [13], influencing choice and attentional allocation in ways similar to explicit reward signals. Stronger sensory evidence biases decisions and accelerates responses in very similar ways as described here [38], and stimuli associated with higher perceptual salience can capture attention similarly to reward-predictive cues [43]. Conversely, distractors associated with reward capture attention more strongly [44]. This suggests that both reward and perceptual information can be integrated within a common framework guiding choice behavior. Yet, such ideas remain to be tested and bounds between costs and rewards may plausibly fall differently, e.g., history-related effects [2] may in some cases shape the priority map similarly to costs.

The differing mappings of costs and rewards onto choice suggest partially dissociable signals prior to utility integration, with candidate sources in noradrenergic, dopaminergic, and cingulate circuits that have been linked to effort and reward processing respectively [25, 26, 45, 46]. The superior colliculus, which receives both inputs and has been implicated in attentional selection and value-based choice [47, 48], is a plausible integration site. These hypotheses are testable through pharmacological or optogenetic manipulation.

We see our framework as a more general and testable extension of existing theories of attentional selection, rather than an alternative. Influential accounts such as the tripartite model [2] have characterized distinct sources of attentional priority, but effort-related costs [and other types of costs, see 20] have been under-theorized, implicitly assumed away, or folded into other constructs rather than treated as an independent determinant of selection. Converging evidence shows that these costs strongly influence attentional choices [5, 15–19, 49, 50] and are directly traded off against rewards. Existing frameworks can thus be understood as specifying the sources of value or priority signals, whereas ours formalizes how these are constrained by costs. The same logic may extend to other value-like signals. Sensory evidence biases decisions and accelerates responses similar to reward [38], perceptually salient stimuli can capture attention much like reward-predictive cues [43], and reward-associated distractors capture attention more strongly than physical salience alone would predict [44]. Reward, salience, and sensory evidence may therefore enter a common utility computation, with costs constraining selection across all of them. This framing also uniquely explains why it is sometimes beneficial to *not* shift attention: when costs exceed rewards for any alternative, staying put is optimal. Converging computational work suggests this efficiency principle operates even deeper, with attentional gain modulation yielding net metabolic savings across the visual hierarchy [51].

One limitation to the linear-sigmoidal dissociation between costs and rewards lies in saccade cost differences being potentially so small that they may only sample the near-linear center of a sigmoid. However, this is precisely the point: for attentional shifts, the extreme zones of a sigmoid would practically never be reached. The operating range of saccade costs is inherently constrained by the geometry of the visual field and the motor system, meaning that a linear function not only accurately describes but may genuinely characterize cost-driven attentional selection under natural conditions. Whether this changes for larger-scale movements - arm, head, or torso - where costs can vary more dramatically, remains a testable open question.

More generally, our results align with frameworks in motor control that have long emphasized fundamental (neuro)economic principles, such as cost minimization and reward maximization [5, 12, 52], but also economic theories, such as theories of attention allocation, where rational inattention formalizes how agents trade information-processing costs against decision utility [7, 8]. For instance, our findings extend the vigor framework [11] from a kinematic readout of utility during deliberation to selection itself. This convergence suggests that movement science, attention research, and behavioral economics may be probing shared computational principles of the brain, albeit through different theoretical lenses and methodological traditions. By integrating costs, rewards and eye movements, the current work is positioned at the intersection between these approaches, and provides empirical support for the notion that theories developed in behavioral economics may provide useful constraints for models of motor control and attentional selection (or vice-versa), highlighting the potential for cross-fertilization of methods. Eye movements are well-suited to study attentional selection and, as we argue, provide an easy-to-study model of economic decision-making. Studying eye-movement behavior may therefore grant researchers in behavioral economics and decision science access to high statistical power and fine-grained measurements of cost-benefit trade-offs. We thus argue that while the scale of decisions may differ across levels, their fundamental working principles are closely aligned.

Finally, we speculate that some phenomena described as irrational in explicit economic decision-making may originate from mechanisms that are adaptive at the level of saccade selection. For instance, the diminishing sensitivity to large reward differences observed here may reflect efficient compression of value signals: once one option is clearly superior, further differentiation yields little additional benefit and would unnecessarily prolong selection, which is consistent with psychometric compression in perceptual decision-making [e.g. 38]. More generally, value representations shaped by attentional efficiency constraints may appear irrational under normative economic models, yet may be well suited for high-frequency, low-level decisions. These links remain speculative and require direct empirical testing, however.

In conclusion, attentional selection follows economic principles: costs and rewards are integrated into a common utility signal that determines eye-movement choices, yet enter it through dissociable computations and scale deliberation differently. By showing that the same cost–benefit logic operating in motor control and value-based choice also governs where we look next, our work establishes attentional selection as a tractable model system for economic decision-making, providing the quantitative cost measurements that earlier utility-based accounts [13, 14] had proposed but could not directly test. Eye movements may thus offer decision science a high-throughput, fine-grained behavioral assay, while economic and movement-science theories offer attention research frameworks in which intrinsic costs sit alongside the priority signals they constrain.

## Methods

### Experiment 1

#### Participants, inclusion, and ethics

Twenty student participants (*M*_age_ = 24.63, *SD*_age_ = 6.25; 14 women, 6 men) with normal or corrected-to-normal vision took part in Experiment 1. Sample size for Experiment 1 was based on effect sizes from Koevoet *et al.* [19], which tested cost and saliency effects on saccade selection with a similar sample size. A priori power analysis indicated 91% power to detect the primary effect of cost on choice (*α* = 0.05). The experimental procedure was approved by Utrecht University’s Faculty of Social Sciences ethical review board (24-0376, 24-0541).

#### Apparatus

Gaze position and pupil size were recorded at 1000 Hz using an Eyelink 1000 desktop mount (SR Research, Ontario, Canada) in a brightness- and sound-attenuated laboratory. A chin- and forehead-rest limited head movements. Stimuli were presented using PsychoPy [v.2024.2.3; 53] on an ASUS ROG PG278Q monitor (2560 x 1440, 100 Hz) positioned 67.5 cm away from eye position. The eye tracker was calibrated and validated (9 points) at the beginning of the session and recalibrated whenever necessary throughout the experiment. The central fixation dot and potential saccade targets were black rings (1°diameter) at an eccentricity of 10°visual diameter presented on a screen-centered circle (12*^◦^* radius, gray ≈42.5 cd/m^2^). The remaining screen was black (≈0.18 cd/m^2^) to ensure equal brightness/contrast adjacent to all target locations.

#### Procedure, task, and stimuli

After providing written informed consent, the experiment started with calibration and validation of the eye tracker. Participants completed a saccade preference task (based on Thomas *et al.* [16] and Koevoet *et al.* [18, 19]; Figure 1a). Importantly, participants were told that three different shapes could be selected (circle, pentagon, rhombus), corresponding to 0 points, 1 point, and 50 points, respectively (values counterbalanced between participants such that circle and rhombus flipped values). This resulted in four reward conditions (equal rewards: 0 and 0, 1 and 1, 50 and 50; 1 vs 0, 50 vs 1, and 50 vs 0) that were present randomly across trials. Points translated into up to €4 extra, in addition to the base €7/hour payment. Participants started trials by looking at the central dot for a random duration between 100-500 ms (within 1.5°of the screen center). Subsequently, two shapes were presented at differing positions at two of the possible, equally spaced 36 target locations (same locations as in Koevoet *et al.* [18]). Participants could then freely select either shape by making an eye movement to the shape. Upon gazing within 1.5°of the center of either shape for more than 100 ms, this shape was counted as selected and participants were fed-back with the associated reward value by black text that was centrally presented.

#### Data processing

All data were processed using custom Python (v3.10) and R (v4.4.3) scripts. Eye-tracking data were downsampled to 250 Hz, and saccades were detected using velocity thresholds as in [18, 19]. For each trial, the selected target was determined based on the final gaze position relative to presented targets. Trials with ambiguous selections or very fast or very slow saccade latencies (<85 ms or >750 ms) were excluded. We used a pupil-derived costs collected in an independent sample wherein participants performed a saccade planning task [data from 18] to obtain effort-costs associated with each position [as in 19].

### Experiment 2

All methods similar to Experiment 1 unless explicitly stated.

#### Participants, inclusion, and ethics

Thirty-two students (*M*_age_ = 22.75, *SD*_age_ = 2.31; 21 women, 11 men) participated in Experiment 2.

#### Procedure, task, and stimuli

Participants first performed a baseline block with two circular shapes, neither of which provided a reward (100 trials per participant). Note that due to an implementation error, not all targets were randomly combined with each other, unlike Experiment 1 and the remainder of Experiment 2. Participants were informed about rewards associated with the three different possible shapes at the start of each trial (see Figure 2a). To this end, they were first presented with the three shapes presented at angular equidistance around the central fixation spot and white text indicating the amount of reward per shape. After a delay of 500 ms, two of these three shapes were presented at different directions. Upon selecting one of the shapes by saccading toward it, the corresponding reward was displayed as feedback. Crucially, shapes were randomly mapped onto directions, such that reward values were associated with directions rather than with the shapes themselves. Thus, reward information for each direction was communicated via the shapes at the beginning of each trial. After feedback, reward values associated with each direction were updated using an ELO-style updating rule [32]; see Figure 2b. Originating from competitive chess rankings, the ELO algorithm updates values based on the discrepancy between the observed choice outcome and the expected outcome derived from the current values. Consequently, updates were smaller when choices were consistent with the model-implied expected preference between the two targets. Participants completed 600 trials each.

#### Data analysis

All statistical tests are two-sided and we set *α* = .05. Direction preferences were analyzed as in [18, Wilkinson notation: *choice* ∼ *obliqueness* + *upness* + *leftness* + (1 + *obliqueness* + *upness* + *leftness* | *experiment*:*participant*)]. Choices were predicted based on differences in costs and rewards between the two saccade targets using a logistic generalized linear mixed-effects model with random intercept and slopes per participant (Wilkinson notation: choice ∼ Δ_cost_ + Δ_reward_ + Δ_cost_:Δ_reward_ + (1 + Δ_cost_ + Δ_reward_ | participant)). Effects of cost and reward differences on saccade latencies were modeled using a linear mixed-effects model, with saccade direction included as a random effect to account for direction-level baseline differences in latency. For conditions with reward differences, we additionally included reward difference as a main effect (Wilkinson notation: onset ∼ Δ_cost_ + Δ_cost_^2^ + [Δ_reward_] + (1 + Δ_cost_ + [Δ_reward_] | participant) + (1 | direction); square brackets denote terms included only when applicable). In two cases, we did not include random slopes because the model failed to converge [54]: Δ_reward_ in Experiment 1 with reward differences, and Δ_cost_ in Experiment 2 BL and Experiment 2. Experiment 2 BL used direction as a fixed covariate rather than a random effect due to limited trials per direction in that block.

## Acknowledgments

We thank Tord Helliesen for assistance with data collection. This project received funding from the Dutch Research Council (NWO), through a Veni grant awarded to Christoph Strauch (VI.Veni.241G.005, 10.61686/EEGGV23807).

## Data availability and code availability

Full materials, data, and analyses are available via the Open Science Framework https://osf.io/cr8gv.

