## Supporting Info for "Attentional selection is a neuroeconomic decision"

The introduction of reward had differing consequences in Experiment 1 and 2. While reward made choices easier in Experiment 1 (i.e., the utility gap becoming wider, reducing choice uncertainty), choices were harder when rewards differed (i.e., the utility gap becoming smaller, increasing uncertainty). If rewards were different, these reward differences did not directly affect latency (unlike cost differences). See Supplementary Figure 1 for a visualization.

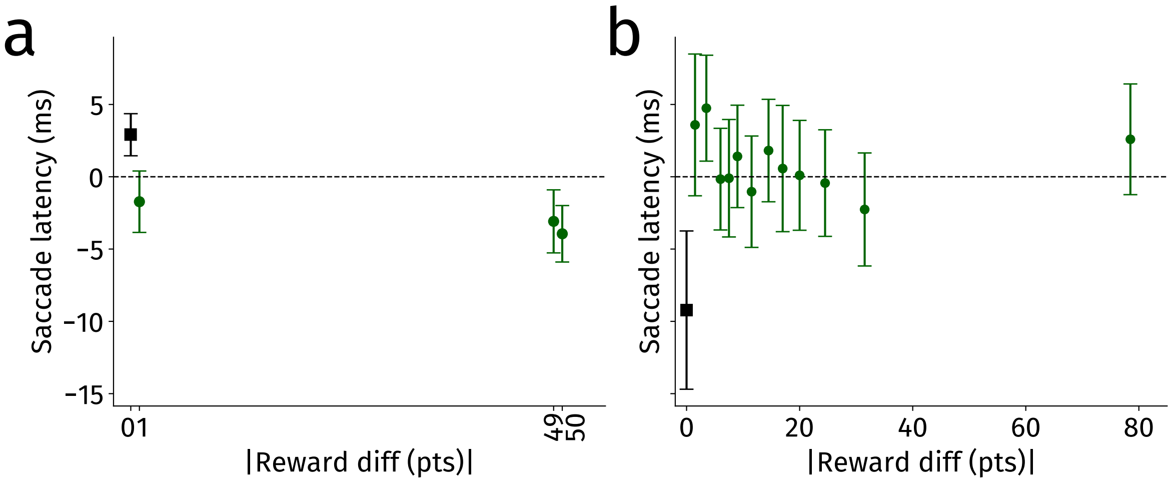


Supplementary Figure 1: Average demeaned (location and participant) saccade latencies in Experiment 1 (a) and Experiment 2 (b). Visualization according to the Figure 5 c,d,e,f for cost differences. Demeaned saccade latencies were not systematically different once reward was present. Saccades were made faster under a reward gap in Experiment 1. When rewards were titrated in Experiment 2, however, this led to slower saccade latencies for the block with a reward difference. The black marker denotes no reward difference between options. Errorbars denote 95% confidence intervals.
